# Human IFITM3 restricts Chikungunya virus and Mayaro virus infection and is susceptible to virus-mediated counteraction

**DOI:** 10.1101/2020.09.11.292946

**Authors:** Sergej Franz, Thomas Zillinger, Fabian Pott, Christiane Schüler, Sandra Dapa, Carlo Fischer, Vânia Passos, Saskia Stenzel, Fangfang Chen, Katinka Döhner, Gunther Hartmann, Beate Sodeik, Frank Pessler, Graham Simmons, Jan Felix Drexler, Christine Goffinet

**Affiliations:** Institute of Experimental Virology, TWINCORE Centre for Experimental and Clinical Infection Research, a joint venture between the Hannover Medical School (MHH) and the Helmholtz Centre for Infection Research (HZI), 30625 Hannover, Germany; Vitalant Research Institute, San Francisco CA, USA; Institute of Clinical Chemistry and Clinical Pharmacology, University Hospital, Venusberg-Campus 1, 53127 Bonn, Germany; Institute of Virology, Campus Charité Mitte, Charité - Universitätsmedizin Berlin, Charitéplatz 1, 10117 Berlin, Germany; Berlin Institute of Health, 10178 Berlin, Germany; Research Group Biomarkers for Infectious Diseases, TWINCORE, Centre for Experimental and Clinical Infection Research, a joint venture between the Hanover Medical School (MHH) and the Helmholtz Centre for Infection Research (HZI), Feodor-Lynen-Str. 7, 30625 Hanover, Germany; Institute of Virology, Hannover Medical School, Carl-Neuberg-Str. 1, 30625 Hanover, Germany; Cluster of Excellence RESIST (EXC 2155), Hannover Medical School, Hannover, Carl-Neuberg-Straße, Germany

**Keywords:** Chikungunya virus/alphavirus, entry restriction, IFITM, viral antagonism

## Abstract

Interferon-induced transmembrane (IFITM) proteins restrict infection by enveloped viruses through interfering with membrane fusion and virion internalisation. The role of IFITM proteins during alphaviral infection of human cells and viral counteraction strategies remain largely unexplored. Here, we characterized the impact of IFITM proteins and variants on entry and spread of Chikungunya virus (CHIKV) and Mayaro virus (MAYV) in human cells, and provide first evidence for a CHIKV-mediated antagonism of IFITM proteins. IFITM1, 2 and 3 restricted infection at the level of alphavirus glycoprotein-mediated entry, both in the context of direct infection and during cell-to-cell transmission. Relocalization of normally endosomal IFITM3 to the plasma membrane resulted in the loss of its antiviral activity. rs12252-C, a naturally occurring variant of *IFITM3* that has been proposed to associate with severe influenza in humans, restricted CHIKV, MAYV and influenza A virus infection as efficiently as wild-type *IFITM3*. Finally, all antivirally active IFITM variants displayed reduced cell surface levels in CHIKV-infected cells involving a posttranscriptional process mediated by one or several non-structural protein(s) of CHIKV.

## INTRODUCTION

Infection of humans by mosquito-transmitted Chikungunya virus (CHIKV) and Mayaro virus (MAYV) causes, in most of the infected individuals, an acute febrile illness accompanied by symmetric joint pain and inflammation. In a subset of infected individuals of varying size depending on the outbreak, long-term morbidity manifesting as chronic arthritis with debilitating pain has been reported for both members of the Semliki Forest Virus complex, and is causing growing medical concern and socio-economical loss in affected countries. While likely being endemic in East Africa since several centuries, Chikungunya virus outbreaks of increasing frequency have occurred worldwide in the past 15 years. MAYV is causing increasing attention in the neotropics. Neither approved vaccines nor antivirals are available against either alphavirus infection. Despite increasing importance of emerging arthritogenic alphaviruses for the human population, the biology of their replication cycle and their interplay with host proteins only begin to be elucidated. Specifically, the entry process of CHIKV and MAYV is greatly facilitated in the presence of target cell-expressed adhesion molecule MXRA8 (Zhang, Kim et al., 2018), and FHL1 serves as a cofactor for CHIKV, but not MAYV, RNA replication (Meertens, Hafirassou et al., 2019).

Interferon-induced transmembrane (IFITM) proteins are broadly active against numerous enveloped RNA viruses, including, among others HIV-1 (Lu, Pan et al., 2011), West Nile virus (Gorman, Poddar et al., 2016), Zika virus (Savidis, Perreira et al., 2016) and influenza A virus (IAV) (Brass, Huang et al., 2009), as well as enveloped DNA viruses (Li, Du et al., 2018, Li, Zheng et al., 2019). Among the five human *IFITM* genes, only *IFITM1*, *IFITM2*, and *IFITM3* have antiviral properties by restricting virus entry (Bailey, Zhong et al., 2014). *IFITM* genes encode small transmembrane proteins of debated topology (Liao, Goraya et al., 2019). IFITM2 and IFITM3 predominantly localize in endosomal membranes, while IFITM1 resides on the cell surface (Chesarino, McMichael et al., 2014, Compton, Roy et al., 2016, Narayana, Helbig et al., 2015, Weston, Czieso et al., 2014). The mechanisms by which IFITM proteins inhibit viral infections appear to involve interference with fusion of viral and cellular membranes, resulting in virions trapped at the hemi-fusion stage (Desai, Marin et al., 2014, Li, Markosyan et al., 2013). IFITM3 retains viral particles in late endosomes and targets them for lysosomal degradation (Feeley, Sims et al., 2011, Spence, He et al., 2019, Suddala, Lee et al., 2019). Experimental retargeting of IFITM3 to the cell surface, which can be induced by disruption of its Yxxθ type endocytosis motif through introduction of the Y^20^A mutation (Williams, Wu et al., 2014) or deletion of the 21 N-terminal amino acids (Jia, Pan et al., 2012, Weidner, Jiang et al., 2010), nullifies its activity against many viruses which enter by endocytosis, including IAV (John, Chin et al., 2013).

The potential importance of human IFITM3 protein as antiviral factor has been addressed in clinical observation studies of Influenza A. Specifically, the SNP rs12252-C allele might associate with increased IAV mortality and morbidity (Everitt, Clare et al., 2012, Pan, Yang et al., 2017, Williams et al., 2014), although other cohorts failed to display this association (Carter, Hebbring et al., 2018, David, Correia et al., 2018, Lopez-Rodriguez, Herrera-Ramos et al., 2016, Mills, Rautanen et al., 2014, Randolph, Yip et al., 2017). A debated mechanistic working model based on the idea that the genetic variation induces alteration of a splice acceptor site, resulting in expression of a truncated IFITM3 protein which lost its antiviral activity (Everitt et al., 2012).

The impact of human IFITM proteins on alphaviral infection remains poorly elucidated, and no information on the anti-alphavirus ability of the SNP rs12252-C allele is available. In *Mus musculus*, a non-natural host of alphaviruses from the Semliki Forest Virus complex, IFITM3 shares 65% homology in amino acid sequence with the human ortholog and inhibits multiple arthritogenic alphaviruses, including CHIKV, and encephalitogenic alphaviruses *in vivo* (Poddar, Hyde et al., 2016). Heterologously expressed human IFITM3, and to a lesser extent IFITM2 restrict Sindbis virus and Semliki Forest virus through inhibition of fusion of viral and cellular membranes (Weston, Czieso et al., 2016). Furthermore, human IFITM1, 2 and 3 emerged as potential inhibitors of CHIKV infection in a high-throughput ISG overexpression screen (Schoggins, Wilson et al., 2011), but their anti-alphaviral properties have not been characterized in the context of CHIKV and MAYV infection.

Here, we addressed the activity of human IFITM1, 2, and 3 as well as of naturally occurring and experimental human IFITM3 variants against CHIKV and MAYV. All three IFITM proteins restricted infection at the level of alphaviral glycoprotein-mediated entry. Experimentally induced relocalization of normally endosomal IFITM3 to the plasma membrane resulted in the loss of its antiviral activity despite robust expression. The rs12252-C allele restricted CHIKV, MAYV and IAV as efficiently as wild-type IFITM3. Finally, steady-state IFITM3 expression levels were markedly reduced at the posttranscriptional level in productively infected cells, suggesting the existence of a so far unappreciated virus-mediated counteraction strategy of IFITM factors mediated by expression of one or several non-structural proteins.

## RESULTS

### Endogenous human IFITM3 restricts CHIKV infection

To investigate the anti-CHIKV activities of human IFITM proteins, we applied a CRISPR/Cas9-assisted approach to edit the *IFITM3* gene in CHIKV-susceptible, IFITM3-expressing HeLa cells (Fig. S1). Specifically, we functionally ablated the *IFITM3* gene by introducing a frameshift after nucleotide 84 (KO) or by deleting a large part of exon 1 of *IFITM3* (Δexon 1) in both alleles. Additionally, we introduced a T-to-C transition at position 89 in both *IFITM3* alleles to express the minor C allele of the single nucleotide polymorphism (SNP) rs12252 (rs12252-C), which has been suggested to associate with severe IAV infections (Everitt et al., 2012, Pan et al., 2017, Zhang, Zhao et al., 2013). Furthermore, we deleted a region of 31 base-pairs encompassing the primary ATG codon (Δ1^st^ ATG) (Suppl. Fig. 1). In all clones, Sanger sequencing confirmed that *IFITM3* (Table 1), but not the highly homologous *IFITM2* gene (data not shown) had been edited. Immunoblotting showed that, while all cell lines and clones clearly upregulated expression of IFITM1 and the prototypic ISG product ISG15 upon IFN stimulation, IFITM3 protein detection was abrogated in HeLa cells encoding *IFITM3* KO and *IFITM3* Δexon1 (Fig. 1A). In addition, no signal was detectable for IFITM3 (Δ1^st^ ATG), arguing against expression of a truncated protein and rather for absence of expression (Fig. 1A). In contrast, three clones expressing the rs12252 T-to-C variant displayed similar IFN-induced IFITM3 expression as parental cells. Importantly, the immunoblot provided no evidence for expression of a truncated IFITM3 protein in rs12252-C cells, but rather displayed a band of equal molecular weight as the one from wild-type IFITM3 (Fig. 1A). Detection of upregulated IFITM2 protein failed despite the confirmed specificity of the IFITM2-targeting antibody (see Fig. 2A). Flow cytometry analysis of IFITM3 protein expression in permeabilized cells paralleled the results of the immunoblot analysis (Figure 1B). Finally, immunofluorescence microscopy confirmed expression of IFITM3 protein in parental and in rs12252-C cells, but not in the other cell lines (Figure 1C).

**Table 1.**
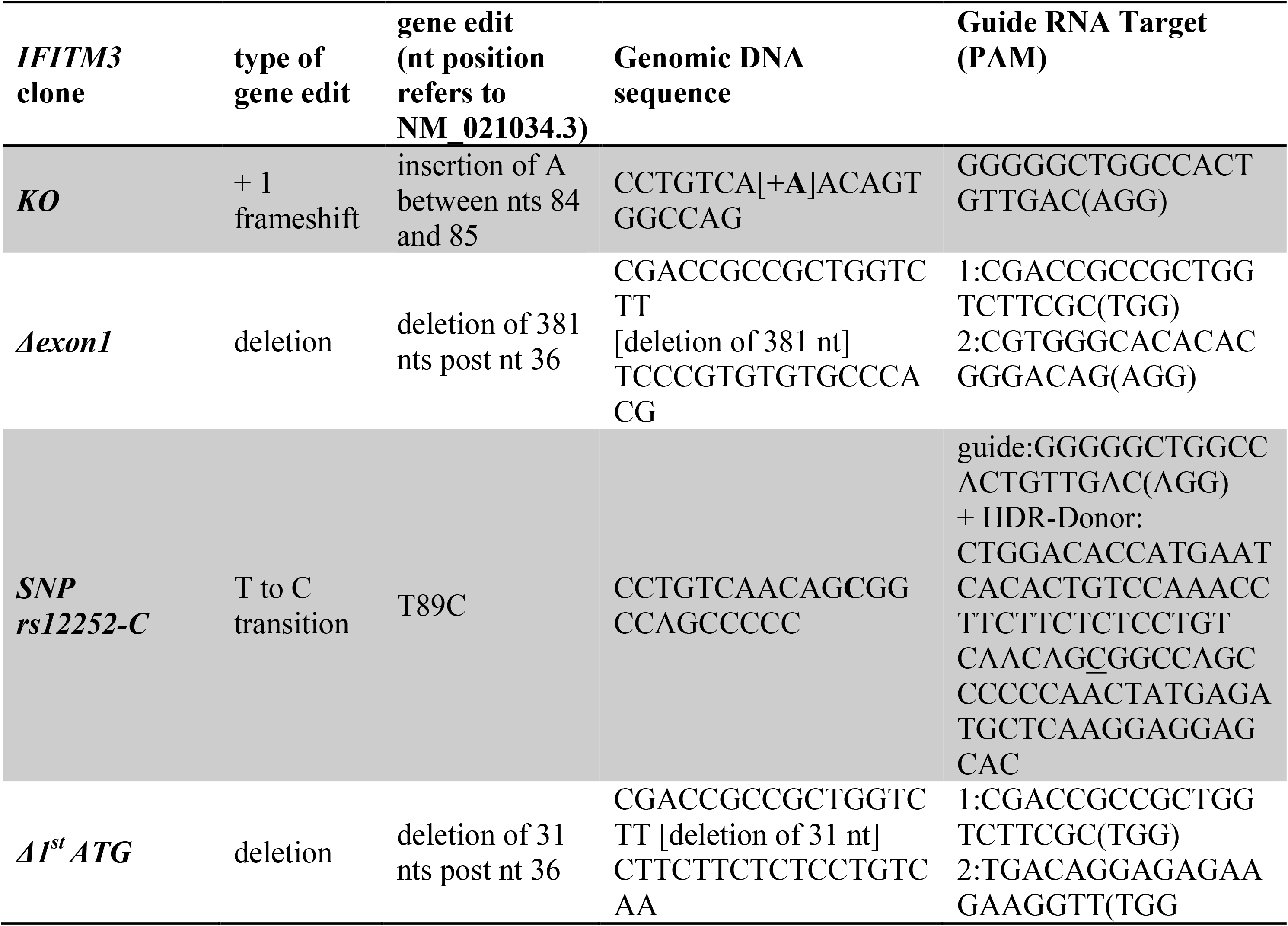
List of *IFITM3* variants generated by gene editing of HeLa cells. nt = nucleotide.

**Figure 1.**
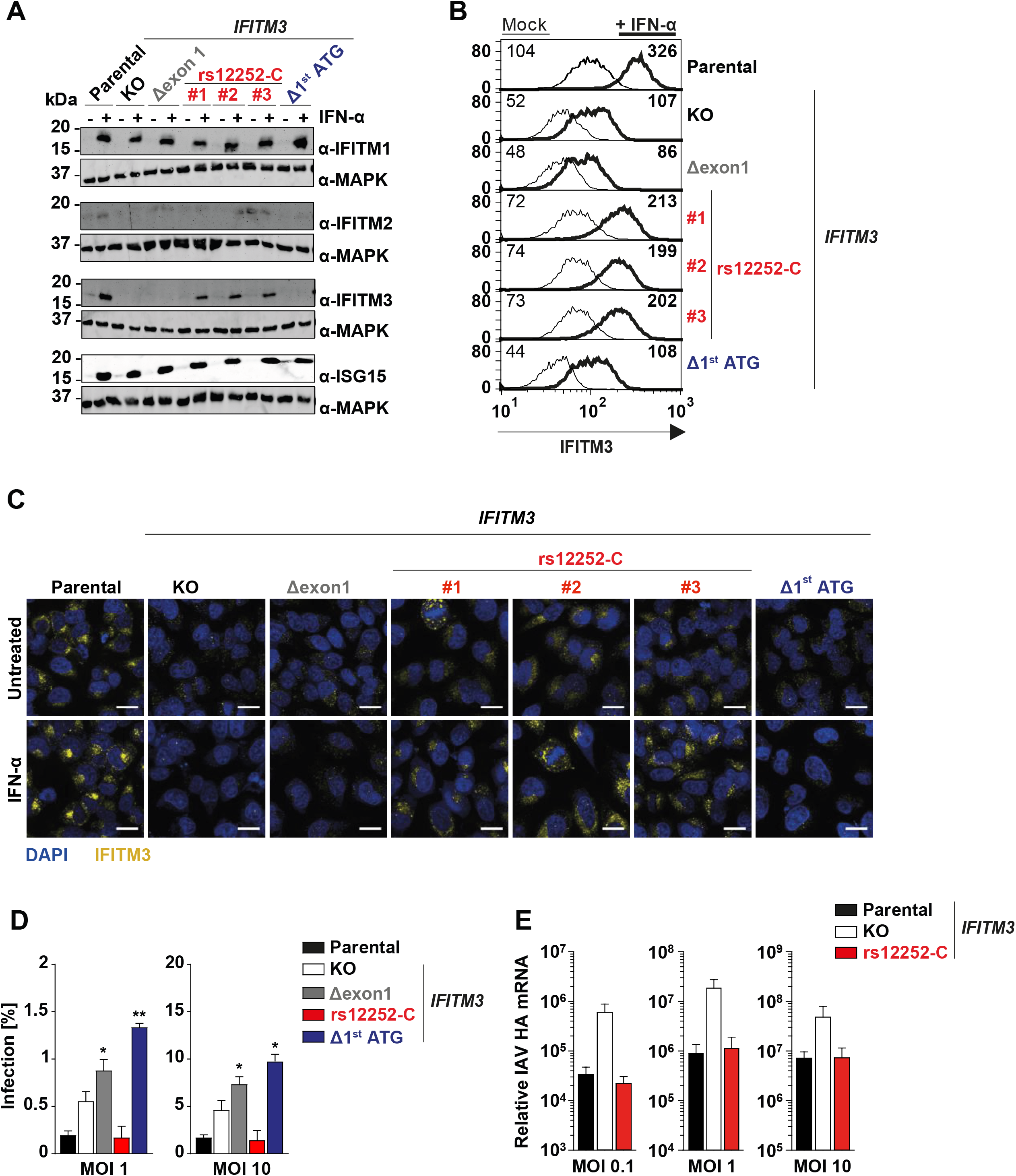
Endogenous human IFITM3 restricts CHIKV infection. (A) Immunoblot of indicated HeLa cell lysates using indicated antibodies. (B) Flow cytometry analysis of IFITM expression in permeabilized HeLa cells. (C) Immunofluorescence microscopy analysis of indicated HeLa cell lines/clones upon mock treatment and treatment with 5000 IU/mL IFN-α for 48 hours (scale bar = 20 μm). (D) HeLa cells were infected with EGFP-CHIKV after a six hours IFN-α pre-treatment duration. The percentage of EGFP-positive infected cells was quantified 24 hours post infection by flow cytometry. (E) HeLa cells were infected with Influenza A virus at indicated MOIs. 24 hours post infection, cell-associated viral HA mRNA was measured by quantitative RT-PCR. SNP: single nucleotide polymorphism, MFI: mean fluorescence intensity, MOI: multiplicity of infection.

**Figure 2.**
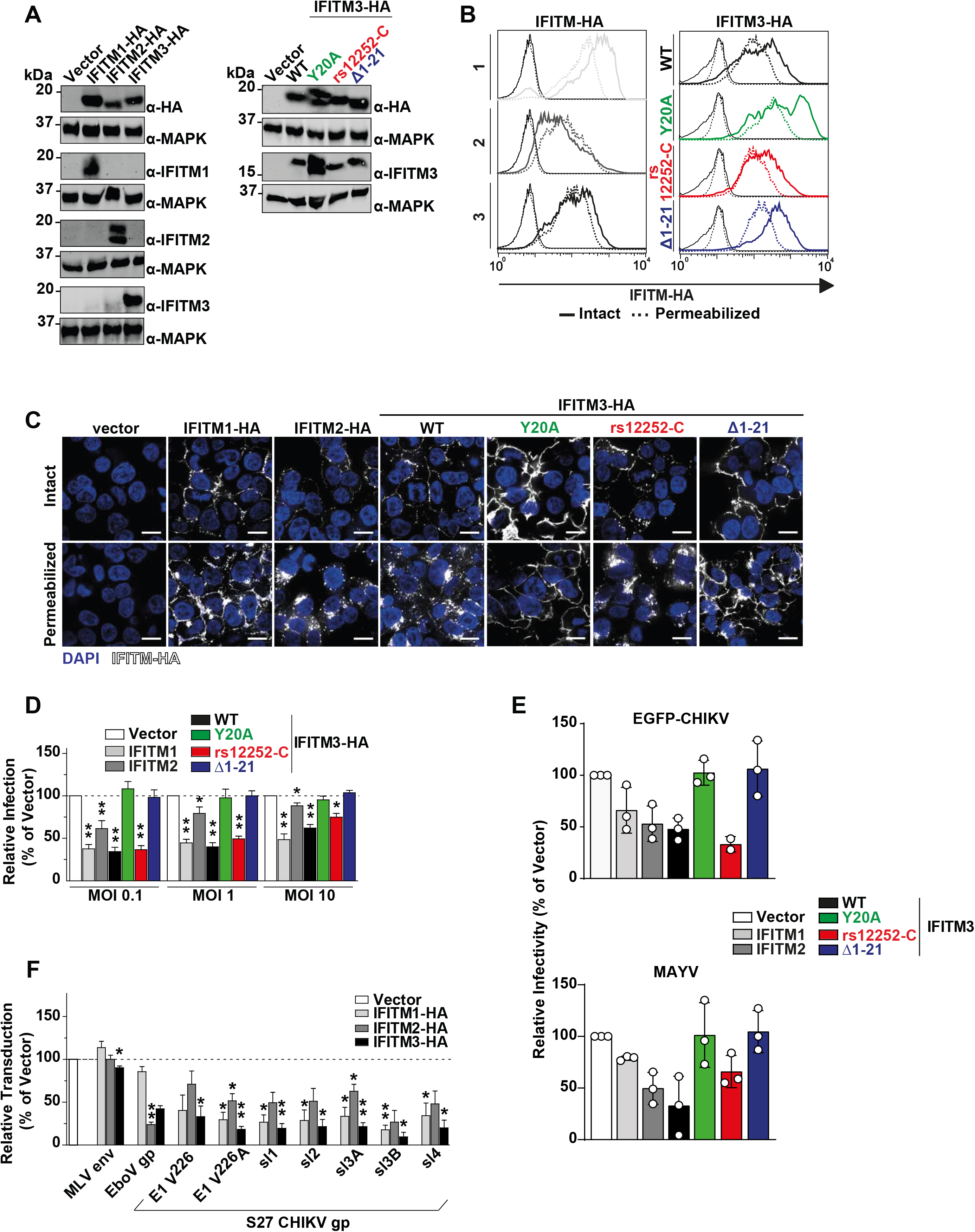
Ectopic expression of human IFITM-HA proteins restricts alphaviral infection by interfering with glycoprotein-mediated entry. HEK293T cell lines stably expressing HA-tagged IFITM proteins were generated via retroviral transduction and puromycin selection. (A) Protein expression was evaluated by immunoblotting using indicated antibodies. (B) Cells were immunostained with an anti-HA antibody and analyzed by flow cytometry, measuring IFITM surface levels in membrane-intact cells and total expression in permeabilized cells. Mean MFI is inset for each condition. (C) Immunofluorescence microscopy of surface and intracellular immunostaining of heterologously expressed IFITM proteins using anti-HA antibody (scale bar = 20 μm). (D) IFITM-HA expressing cells were infected with EGFP-CHIKV of increasing MOIs for 24 hours, and the percentage of EGFP-positive cells was quantified by flow cytometry. (E) Stable cell lines were inoculated for 48 hours with lentiviral firefly luciferase-expressing reporter pseudoparticles decorated with glycoproteins of murine leukemia virus, Ebola virus, or different CHIKV lineages. Luciferase activity in transduced cells was quantified luminometrically and normalized to the vector control cell line.

We then inoculated the individual cell lines with CHIKV. Throughout the study, we used the Indian Ocean lineage strain LR2006 harboring an EGFP reporter at the 5’ end of the viral genome, hereafter referred to as EGFP-CHIKV. Absence of IFITM3 expression in the KO, Δexon1 and Δ^1st^ ATG cells was accompanied by diminished effectivity of the IFN-induced antiviral program, while IFITM3 expression in parental and rs12252-C cells impaired EGFP-CHIKV infection to similar levels (Fig. 1D). Finally, parental cells and rs12252-C cells were equally restrictive to IAV infection, as opposed to KO cells (Fig. 1F). In conclusion, we show that endogenous IFITM3 restricts CHIKV infection. Furthermore, the naturally occurring variant of *IFITM3*, rs12252-C, drove expression of an IFITM3 protein that displays a similar anti-CHIKV and anti-IAV potency as the major *IFITM3* allele.

### Ectopic expression of human IFITM-HA proteins restricts alphaviral infection by interfering with glycoprotein-mediated entry

To gain mechanistic insight into the differential antiviral potency of IFITM proteins and variants, we stably transduced HEK293T cells, which lack endogenous IFITM protein expression, with C-terminally HA-tagged IFITM1, 2, or 3 (Brass et al., 2009). In addition, we generated individual cell lines expressing the mutant proteins IFITM3(Y^20^A)-HA and IFITM3(Δ1-21)-HA, which localize to the plasma membrane (Feeley et al., 2011, Jia et al., 2012, Weidner et al., 2010, Williams et al., 2014). Finally, we overexpressed the IFITM3 rs12252-C variant (IFITM3(rs12252-C)-HA). Appropriate protein expression was confirmed by immunoblot analysis using both an anti-HA antibody and primary antibodies targeting the individual authentic IFITM proteins (Fig. 2A). We confirmed similar expression levels of all proteins and variants by flow cytometry (Fig. 2B) and immunofluorescence microscopy (Fig. 2C) using an anti-HA antibody in intact and in permeabilized cells. Thereby, we confirmed the previously reported, predominant cell surface localization of IFITM3(Y^20^A)-HA and IFITM3(Δ1-21)-HA (Perreira, Chin et al., 2013). IFITM3(rs12252-C)-HA did not express a shorter variant and the protein localized predominantly intracellularly, as observed in HeLa cells (Fig. 1). To assess the antiviral capacity of the IFITM-HA proteins, we infected the cell lines with EGFP-CHIKV and determined the amount of EGFP-positive cells at 24 hours post infection. Over a wide range of MOIs, IFITM1-HA and IFITM3-HA, and to a lesser extent IFITM2-HA, displayed antiviral activity against CHIKV (Fig. 2D, Supplemental Fig. S2). IFITM3(rs12252-C)-HA restricted EGFP-CHIKV infection at a similar efficiency as IFITM3-HA. In contrast, IFITM3(Y^20^A)-HA and IFITM3(Δ1-21)-HA failed to impair infection (Fig. 2D, Supplemental Fig. S2). We observed a similar pattern of inhibition at the level of infectivity released in the culture supernatant of EGFP-CHIKV- and MAYV-infected cells, as judged by plaque assays (Fig. 2E). To monitor whether inhibition of CHIKV infection was related to the established ability of IFITMs to interfere with enveloped virus entry, we quantified the cells’susceptibility to transduction by lentiviral pseudoparticles decorated with heterologous viral glycoproteins and expressing a luciferase cassette (Fig. 2F). In line with previous reports (Brass et al., 2009), IFITM protein expression did not reduce the susceptibility of cells to transduction by particles pseudotyped with glycoproteins of murine leukemia virus. In contrast, expression of IFITM2-HA and IFITM3-HA, but not IFITM1-HA, reduced the cells’ susceptibility to transduction by Ebola virus glycoprotein-pseudotyped particles, as reported before (Brass et al., 2009, Wrensch, Karsten et al., 2015). Transduction of cells by particles incorporating CHIKV glycoproteins was impaired by IFITM1-HA and IFITM3-HA and, to a lesser extent, by IFITM2-HA. The A^226^V mutation in CHIKV E1 that enabled adaptation of the virus to the alternative vector *Aedes albopictus* (Tsetsarkin, Vanlandingham et al., 2007) did not overtly modulate susceptibility to IFITM protein-mediated restriction (Fig. 2F). CHIKV glycoproteins from sublineages 1-4, which are derived from the S27 strain (Tsetsarkin, Chen et al., 2014), shared susceptibility to IFITM-mediated inhibition, although to slightly different extents (Fig. 2F). Collectively, these data establish that IFITM proteins restrict CHIKV and MAYV, and that inhibition is directed against entry mediated by glycoproteins of different CHIKV sublineages. Additionally, antiviral activity is only exerted by IFITM3 variants with reported endosomal localization.

### Cell-to-cell transmission of and direct infection by CHIKV differ in efficiency, but share susceptibility to IFITM-mediated restriction

Cell-to-cell transmission of virions can antagonize or evade the IFN-induced antiviral state, including restriction by some (Jolly, Booth et al., 2010, Richardson, Carroll et al., 2008, Vendrame, Sourisseau et al., 2009), but not all (Puigdomenech, Casartelli et al., 2013) antiviral factors. Expression of IFITM3 renders target cells less susceptible to infection by cell-free HIV-1 particles, but cell-to-cell transmission of HIV-1 partially escapes this antiviral effect (Compton, Bruel et al., 2014). We thus addressed the relative ability of cell-associated and cell-free transmission modes in supporting CHIKV spread, and their relative susceptibility to IFITM protein-mediated restriction. As a surrogate system for cell-to-cell transmission, we subjected infected cells with an agarose overlay, which favors transfer of virions between neighboring cells and impairs infection by freely diffusing virus particles. In this set-up, the initial infection is seeded by cell-free virus, whereas subsequent rounds of infection are mediated by cell-associated virus and cell-to-cell spread. For comparison, infected cells were cultured in regular growth medium without agarose. EGFP-CHIKV spread was enhanced under agarose overlay conditions, with to two-fold higher percentages of EGFP-positive cells as compared to cells cultured in regular medium (Fig. 3A). Of note, in cells under agarose, an EGFP-encoding HSV-1 spread at identical or even lower efficiencies than in regular medium (Fig. 3A). Fluorescence microscopy analysis of EGFP-CHIKV-infected cells cultured under agarose overlay revealed a nest-like pattern, reflecting spread of one initial infection event to closely surrounding cells through intercellular contacts, in contrast to the more evenly distributed, punctuated pattern consistent with spread of the infection by freely diffusing virus particles (Fig. 3B).

**Figure 3.**
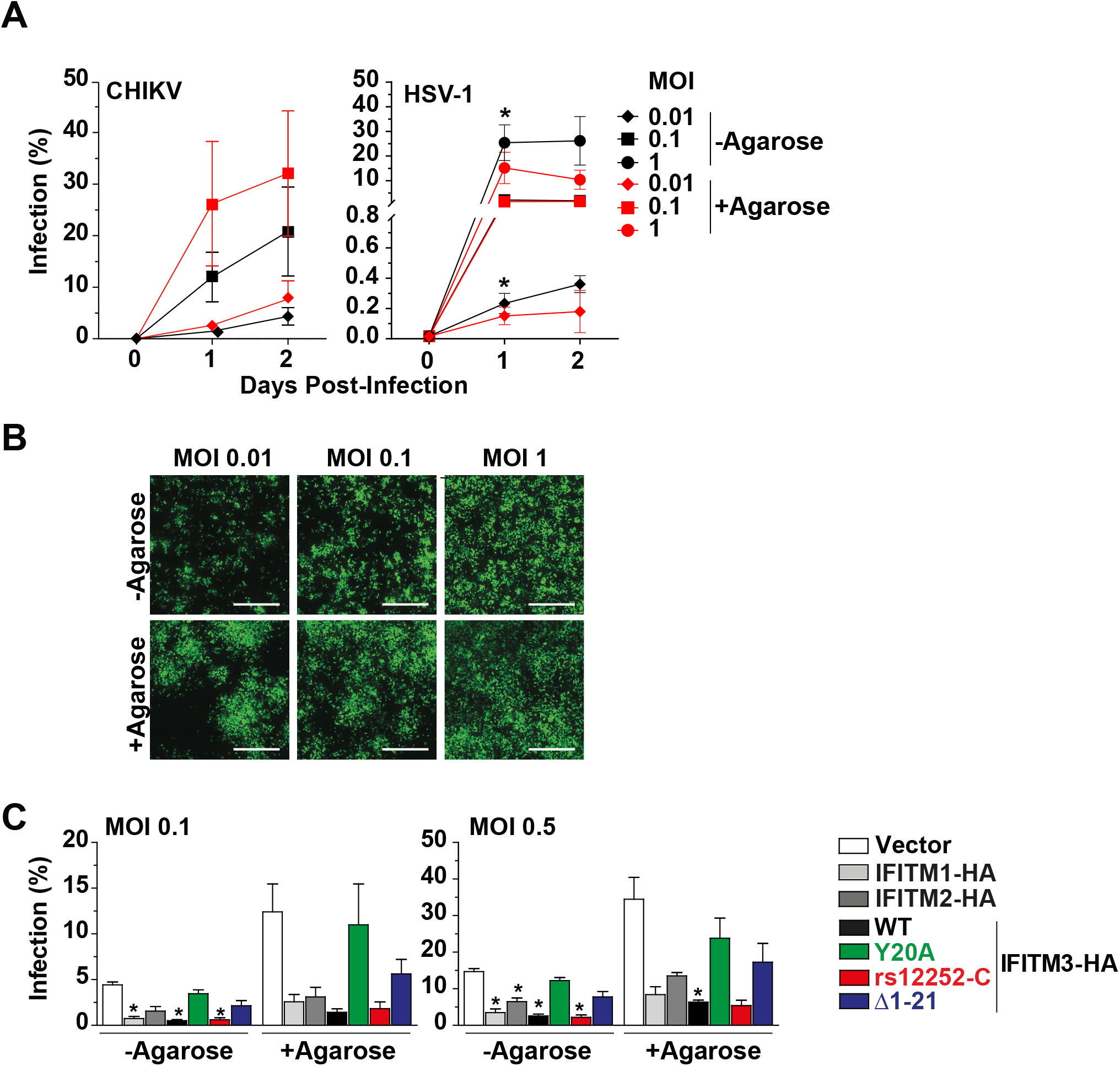
Cell-to-cell transmission of and direct infection by CHIKV differ in efficiency, but share susceptibility to IFITM-mediated restriction. Confluent HEK293T cells were infected with EGFP-CHIKV or EGFP-HSV-1 for six hours. Subsequently, cells were overlaid with 0.8 % agarose or left untreated. (A) EGFP-positive cells were measured one and two days post infection via flow cytometry and (B) microscopic analysis of CHIKV-infected cells was performed 24 hours post infection (scale bar = 500 μm). (C) Confluent HEK293T cells ectopically expressing IFITM-HA proteins were infected with EGFP-CHIKV for six hours followed by the application of a 0.8 % agarose overlay. 24 hours post infection, EGFP-positive cells were measured via flow cytometry.

We next investigated the restriction potential of the individual IFITM proteins and variants in the context of cell-free and cell-associated CHIKV infection (Fig. 3C). For any cell line, the percentage of EGFP-positive cells was consistently elevated under agarose overlay conditions as compared to culture in regular medium. The fold increase was similar in all cell lines, indicating that agarose overlay treatment yielded a better spread irrespective of expression of IFITM proteins. Importantly, CHIKV was unable to escape IFITM-mediated restriction by cell-to-cell transmission (Fig. 3C). Taken together, CHIKV spreads efficiently through cell-to-cell transmission, surpassing the spreading efficiency in a regular culture in which virions diffuse freely. However, the restriction pattern of IFITM proteins and variants is maintained in the context of cell-to-cell transmission.

### Reduction of cell surface expression of antiviral IFITM proteins in CHIKV-infected cells

Flow cytometric analysis of CHIKV-infected cells showed a markedly lowered cell surface abundance of most, but not all IFITM-HA proteins in productively infected, EGFP-positive cells as opposed to bystander, EGFP-negative cells of the identical culture (Fig. 4A). The decrease of cell surface levels was most pronounced for IFITM3-HA and IFITM-3(rs12252-C)-HA, followed by IFITM2-HA. Surface levels of IFITM1-HA, IFITM3(Y20A)-HA, and IFITM3(Δ1-21)-HA, which share a predominantly cell surface localization, were clearly less affected (Fig. 4A-B). We confirmed this finding by immunofluorescence microscopy (Fig. 4C). To address if surface reduction of selected IFITM-HA proteins in EGFP-positive cells requires the entry process of CHIKV infection, we bypassed the entry step by directly transfecting the full-length CHIKV RNA genome. Strikingly, IFITM-HA proteins were also downregulated in EGFP-CHIKV-transfected cells, in stark contrast to cells transfected with HIV-1 EGFP DNA (Fig. 4D). Furthermore, HEK293T IFITM3-HA cells transfected with either EGFP-CHIKV genomic RNA or mRNA encoding CHIKV nsP1-4 and EGFP shared reduction of IFITM3-HA levels on their surface specifically in EGFP-positive cells (Fig. 4E). These data suggest that expression of one or several nonstructural CHIKV proteins induces the downregulation of antivirally active IFITM proteins. In contrast, inactive IFITM3 variants that localize predominantly at the cell surface are less susceptible to this downregulation.

**Figure 4.**
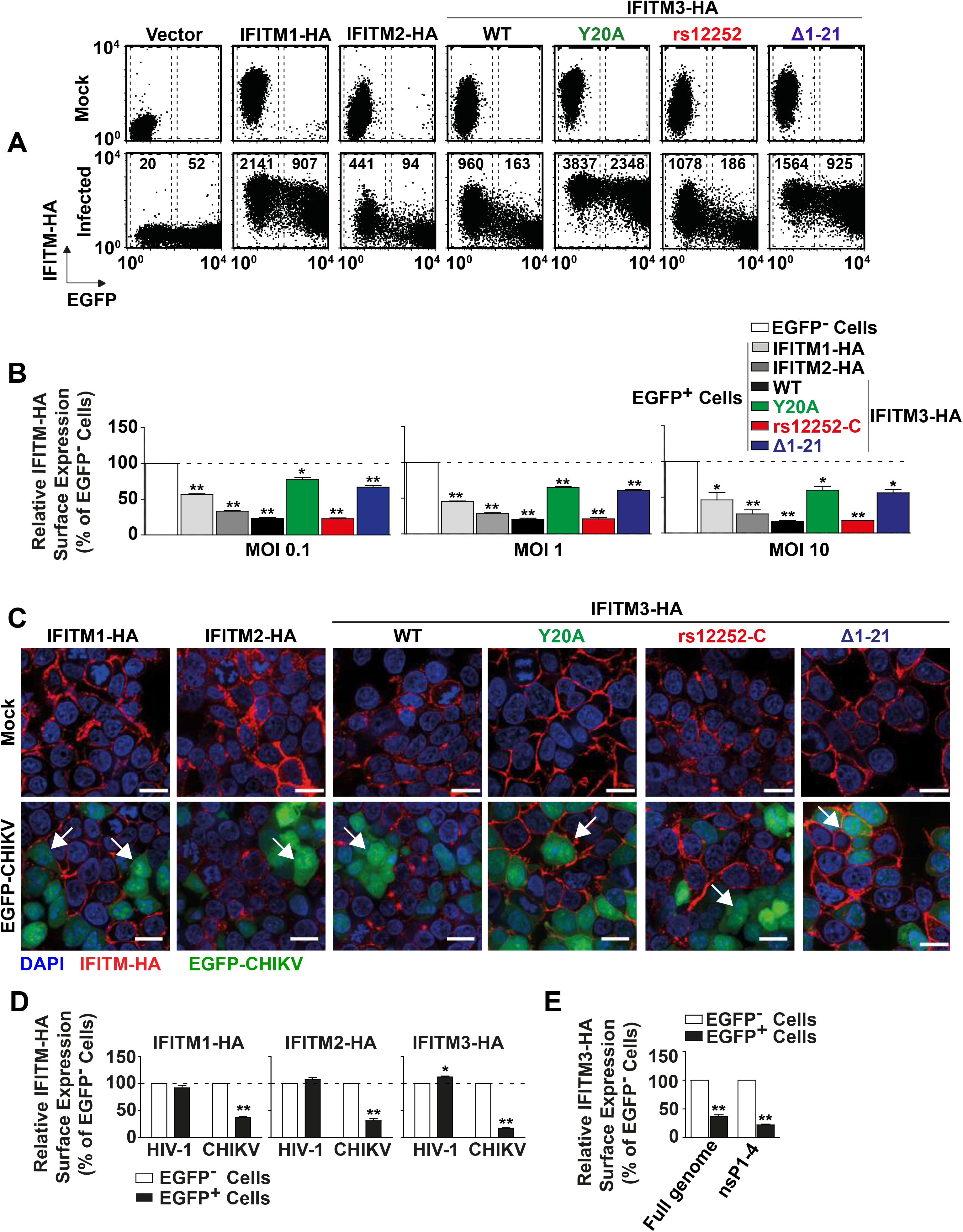
Reduction of cell surface expression of antiviral IFITM proteins in CHIKV-infected cells. (A) Indicated HEK293T cells were infected with EGFP-CHIKV (MOI 10) for 24 hours and were immunostained with an anti-HA antibody for flow cytometry. (B) Quantification of IFITM-HA expression in EGFP-positive relative to EGFP-negative cells after infection with EGFP-CHIKV for 24 hours. (C) Indicated HEK293T cell lines were infected with EGFP-CHIKV (MOI 10) for 24 hours. Permeabilized cells were immunostained with an anti-HA antibody and analyzed microscopically (scale bar = 20 μm). Arrowheads indicate EGFP-positive cells with IFITM-HA expression (red) or lack thereof. (D) Indicated cell lines were transfected individually with full-length EGFP-CHIKV RNA and HIV-1 EGFP DNA and were immunostained with an anti-HA antibody. MFIs of IFITM-HA EGFP-positive cells were determined via flow cytometry and normalized to the MFI of EGFP-negative cells. (E) Quantification of IFITM3-HA surface level 24 h after individual transfection of HEK293T IFITM-3-HA cells with full length EGFP-CHIKV mRNA or RNA encoding the non-structural proteins 1-4 and EGFP.

### Posttranscriptional reduction of endogenous IFITM3 expression in CHIKV-infected cells

Next, we analyzed the consequence of CHIKV infection on endogenous IFITM3 expression levels. By immunofluorescence microscopy, we quantified the amounts of IFITM3 protein in productively infected, EGFP-positive and bystander, EGFP-negative HeLa cells (Fig. 5A). Similar to the heterologous expression set-up (Fig. 4), we detected a strong reduction of endogenous IFITM3 immunostaining intensity in highly EGFP-positive cells, while infected cells with low EGFP intensity and EGFP-negative cells displayed similarly high levels of IFITM3 (Fig. 5 A-B). In stark contrast, ISG15 and MX1 protein levels remained similar in EGFP-positive and EGFP-negative cells (Fig. 5C), arguing against a global translational shut-off as the underlying reason and pointing towards a specific effect on IFITM3. Finally, CHIKV infection failed to modulate *IFITM3* mRNA expression in HeLa cells (Fig. 5D), excluding that infection modulates transcription of the *IFITM3* gene, and suggesting an infection-induced modulation of IFITM3 protein steady-state levels. IFN-α treatment boosted *IFITM3* mRNA levels, as expected. Furthermore, another prototypic ISG, *IFIT1*, was clearly induced by both IFN-α treatment and CHIKV infection (Fig. 5E). Conclusively, these data support a posttranscriptional modulation of IFITM3 expression in CHIKV-infected cells, potentially representing a virus-mediated counteraction strategy of IFITM restriction.

**Figure 5.**
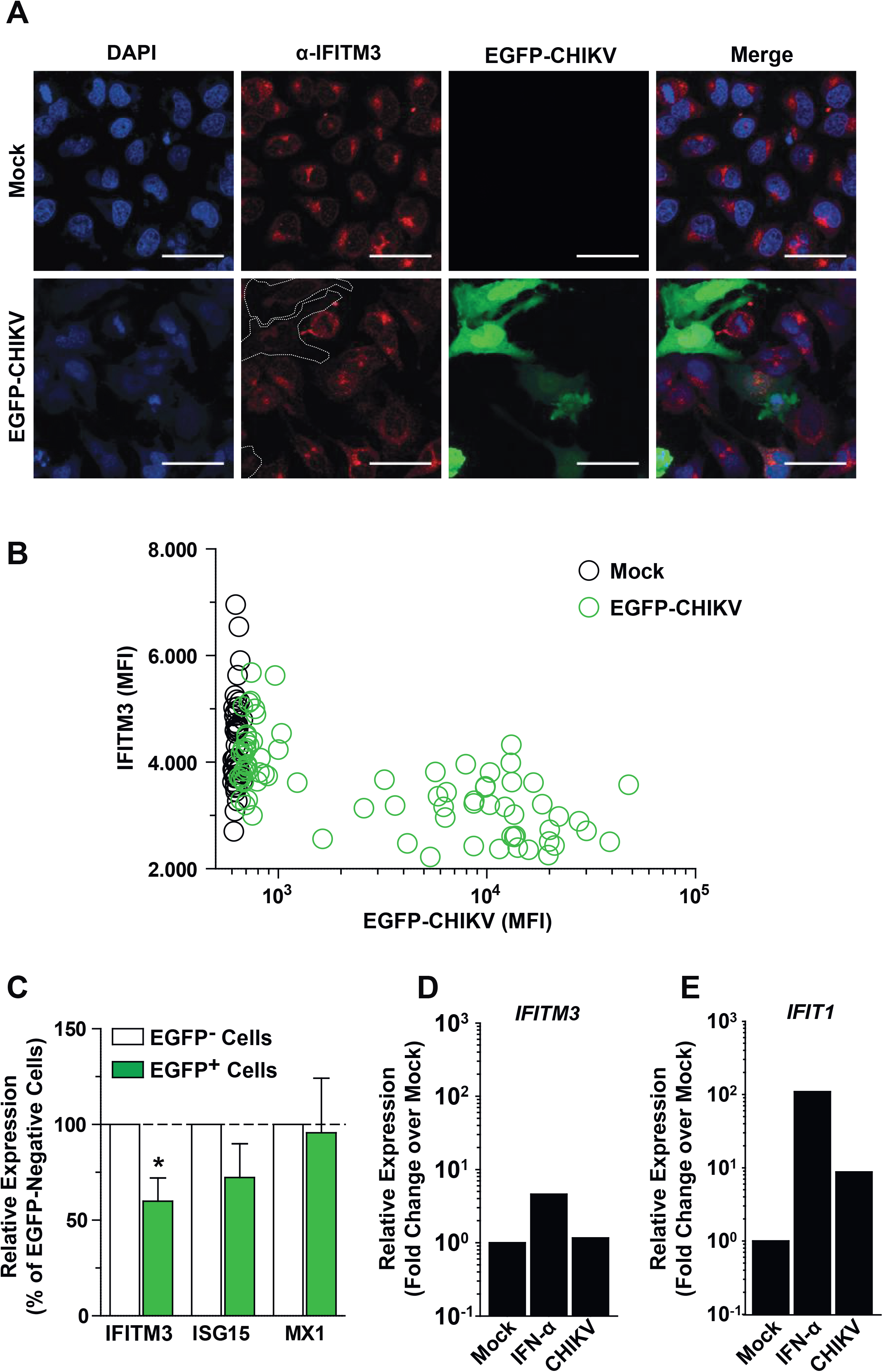
Posttranscriptional reduction of endogenous IFITM3 expression in CHIKV-infected cells. (A) HeLa cells were infected with EGFP-CHIKV (MOI 100) for 24 h, permeabilized, and immunostained for IFITM3 prior to microscopic analysis. Dotted lines indicate the border of EGFP-positive cells (scale bar = 50 μm). (B) Quantification of microscopic images of infected HeLa cells. IFITM3 MFI was determined using ImageJ and is plotted against EGFP MFI (mock cells n = 50; CHIKV infected cells n = 80). (C) HeLa cells were infected with EGFP-CHIKV (MOI 10) for 24h. Subsequently, permeabilized cells were stained for IFITM3, ISG15 or MX1 and MFI values were normalized to EGFP-negative cells. (D) HeLa cells were infected with EGFP-CHIKV and 24 hours post infection, *IFITM3* and *IFIT1* mRNA levels were measured by quantitative RT-PCR.

## DISCUSSION

First evidence for anti-CHIKV properties of human IFITM proteins 1, 2 and 3 was provided in a high-throughput ISG overexpression screen in STAT-deficient human fibroblasts (Schoggins et al., 2011). However, the pattern of IFITM-mediated CHIKV inhibition and potential CHIKV-mediated counteraction strategies remain obscure. By applying a two-armed approach that included investigation of endogenous *IFITM3* and variations thereof, and of heterologously expressed, epitope-tagged IFITM proteins and mutants, we established and characterized the antiviral activity of IFITM proteins against CHIKV and MAYV.

Our study centered on the investigation of IFITM3’s role on alphaviral infection. Expression of endogenous and heterologous IFITM3 rendered HeLa and HEK293T cells less susceptible to CHIKV infection, respectively. In the endogenous expression context, frameshift insertion in the 5’part of the *IFITM3* gene, deletion of a large part of the first exon of the *IFITM3* gene or of a smaller, 31 base-pair region comprising the first, canonical ATG, resulted in ablation of IFITM3 expression and antiviral activity. Similarly, cells expressing a frameshift in the 5’ part of the first exon of the *IFITM3* gene were unable to restrict IAV infection, and lost the ability to restrict CHIKV.

It has been hypothesized that the T-to-C substitution in the SNP rs-12252, located in the first exon of the *IFITM3* gene, alters a splice acceptor site, resulting in a truncated IFITM3 protein lacking its first 21 amino acids and exerting reduced antiviral activity (Everitt et al., 2012). However, this working model did not substantiate, since neither the predicted alternatively spliced mRNA (Makvandi-Nejad, Laurenson-Schafer et al., 2018, Randolph et al., 2017) nor a truncated IFITM3 protein (Makvandi-Nejad et al., 2018) has been detected in cells homozygously expressing rs12252-C. In line with these negative results, we detect a protein of normal size both in cells homozygously expressing the SNP and in cells overexpressing an IFITM3-encoding construct carrying the T-to-C transition. However, our anti-IFITM3 immunoblotting technique is clearly able to detect the smaller molecular weight of IFITM3(Δ1-21)-HA, excluding a technical inability to detect marginally smaller IFITM3 proteins in general. While IFITM3(Δ1-21)-HA localized to the cell surface and lacks antiviral activity, subcellular localization and antiviral phenotype of the rs12252-C variant remained indistinguishable from the wild-type IFITM3 in terms of anti-CHIKV, anti-MAYV and anti-IAV-activity, molecular weight and subcellular localization. Together, we conclude that if any, specific functional properties of the rs12252-C allele remain to be discovered and may not directly be implicated in cell-intrinsic immunity.

Experimental redirection of IFITM3 to the cell surface by disrupting the sorting motif YxxФ (Jia et al., 2012, Jia, Xu et al., 2014) through introduction of the Y^20^A or Δ1-21 mutations resulted in a loss of its anti-CHIKV and anti-MAYV activity. Analogous findings have been obtained by others for SINV and SFV (Weston et al., 2016), and other enveloped viruses which invade cells via receptor-mediated endocytosis (Jia et al., 2014, John et al., 2013). In addition, heterologous assays revealed an anti-CHIKV and anti-MAYV activity of IFITM1 that equaled that of IFITM3. This observation contrasts reports by others for Sindbis virus and Semliki Forest virus (Weston et al., 2016), but is in accordance with results obtained in a study that screened the antiviral potential of several ISGs against CHIKV (Schoggins et al., 2011). Restriction was directed against CHIKV E2/E1 glycoprotein-mediated entry, was maintained in the context of CHIKV glycoproteins expressing the vector switch-enabling mutation A^226^V in CHIKV E1, and displayed a significant breadth since glycoproteins of several CHIKV S27 strain sublineages were sensitive to IFITM protein-mediated restriction.

The majority of cell culture studies on CHIKV is conducted using cell-free, purified virus. An early report published in 1970 suggested that CHIKV spreads via cell-to-cell contacts when free virions are immunologically neutralized (Hahon & Zimmerman, 1970). Along this line, intercellular transmission of CHIKV was reported more recently to be less sensitive to antibody-mediated neutralization (Lee, Kam et al., 2011). In the present study, we applied a semi-solid, agarose-containing overlay to infected cultures to determine the contribution of cell-free virus versus intercellular transmission, as performed by others for hepatitis C virus (Timpe, Stamataki et al., 2008), vesicular stomatitis virus and murine leukemia virus (Liberatore, Mastrocola et al., 2017). Interestingly, CHIKV tended to spread more efficiently under agarose overlay, as opposed to HSV-1. The exact mode of intercellular transmission of CHIKV in different cell types will be an important future object of investigation and might contribute to understanding alphavirus persistence *in vivo*. The relative restriction potential of individual IFITM proteins and variants was identical in the cell-free and cell-associated transmission set-ups. As a contrasting example, HIV-1 is able to overcome IFITM3-mediated restriction via cell-to-cell spread, however only when IFITM3 is expressed solely on target cells (Compton et al., 2014). Remarkably, IFITM proteins display a second layer of antiviral activity, which consists in diminishing the infectivity and fusogenicity of enveloped virus particles produced in IFITM protein-expressing cells (Compton et al., 2014, Tartour, Appourchaux et al., 2014). This process is often, but not always, accompanied by incorporation of IFITM proteins into the membrane of secreted virions (Appourchaux, Delpeuch et al., 2019). Future studies are required to characterize susceptibility of CHIKV particles to IFITM-mediated negative imprinting and whether the mode of viral counteraction identified in this study is related to this site of action rather than to entry inhibition.

CHIKV has evolved several strategies to evade or antagonize cell-intrinsic immunity, facilitating successful replication and spread in its host. The multifunctional nonstructural protein nsP2 of CHIKV, besides proteolytically processing the nonstructural polyprotein precursor (Rausalu, Utt et al., 2016), inhibits IFN-induced nuclear translocation of STAT1 in Vero cell lines (Fros, Liu et al., 2010). Furthermore, it degrades the Rpb1 subunit of the RNA polymerase II in BHK-21 and NIH3T3, but not mosquito cells (Akhrymuk, Kulemzin et al., 2012). Additionally, translational shutoff has been observed in some CHIKV-infected cell lines (White, Sali et al., 2011). Here, in two different cellular systems, we obtained no evidence for a broad transcriptional or translational shutoff indicating some cell line or cell type specificity. On the contrary, productive CHIKV infection associates with reduced IFITM3 protein levels, and this reduction appears to operate at the posttranscriptional level. In the HEK293T-based heterologous expression system, only antivirally active, endosomally located (WT; rs-12252-C) but not inactive, plasma membrane-resident (Y^20^A; Δ1-21) IFITM3 proteins were reduced in quantity in CHIKV-infected cells. This specificity was observed despite all IFITM-HA proteins being heterologously expressed under the control of the identical CMV immediate early promoter, which is unlikely to be targeted by CHIKV evasion strategies. In CHIKV-infected HeLa cells, endogenous IFITM3 protein levels were reduced in the absence of a detectable net decrease of *IFITM3* mRNA, again pointing towards a specific counteraction strategy directed against the IFITM3 protein. Interestingly, IFITM3 degradation has been reported to occur through ubiquitination of a highly conserved PPxY motif that overlaps with the aforementioned YxxФ endocytosis motif by the E3 ubiquitin ligase NEDD4 (Chesarino, McMichael et al., 2015). With Y^20^ representing both a critical phosphorylation site required for IFITM3 internalization and part of a ubiquitination motif important for degradation of the protein (Chesarino et al., 2014, Yount, Karssemeijer et al., 2012), it is tempting to speculate that a nonstructural protein of CHIKV promotes, either directly or indirectly, endocytosis and/or ubiquitination-dependent degradation of IFITM3, a process that has yet to be studied in detail.

## MATERIALS & METHODS

### Cell lines

BHK-21, HeLa (ATCC® CCL-2) and HEK293T (ATCC® CRL-3216) cells were cultured in Dulbecco’s modified Eagle’s medium (DMEM) with 10 % fetal bovine serum (FBS), 100 μg/mL streptomycin, and 2 mM L-glutamine. For an agarose-overlay, pre-warmed DMEM with 2 % FBS mixed with liquid SeaPlaque™ Agarose (Lonza) to a final concentration of 0.8 % agarose was added to cells two hours post infection. HEK293T cells expressing vector or IFITM-HA proteins were generated via retroviral transduction and subsequent puromycin selection. Interferon stimulation was performed using indicated concentrations of Roferon (Interferon-α2a, Roche).

### Gene editing

*IFITM3*-edited HeLa clones were generated by electroporation of pMAX-CRISPR plasmids encoding EF1α promotor-driven Cas9-2A-EGFP and U6 promoter-driven chimeric guide RNAs (gRNAs) via the Neon Transfection System (Thermo Fisher Scientific, Darmstadt, Germany), settings were as recommended by the manufacturer (Neon cell line database, Hela cells): 5×10^6^ cells/ml, 1005 V, 35 ms pulse width, 2 pulses. 50 μg/ml of each plasmid was used, for generation of *rs12252* point mutation via HDR, 5μM ssDNA repair template (Integrated DNA Technologies, Leuven, Belgium) was added. After electroporation, EGFP-expressing cells were FACS-sorted and single cell clones were obtained by limiting dilution. For gRNA target sequences and HDR-template sequence see Table 1. PCR-amplified gene loci of individual clones were genotyped by Sanger sequencing (SeqLab, Göttingen, Germany) using the following primers. IFITM3 locus forward: TTTGTTCCGCCCTCATCTGG; IFITM3 locus reverse1 (KO, rs12252. Δ1stATG): CACCCTCTGAGCATTCCCTG, IFITM3 locus reverse2 (Δexon1): GTGCCAGTCTGGAAGGTGAA, IFITM2 locus forward: CCCTGGCCAGCTCTGCA and IFITM2 locus reverse: CCCCTGGATTTCCGCCAG.

### Plasmids, Retro- and Lentiviral Vectors

Individual pQCXIP constructs encoding IFITM1-HA, IFITM2-HA and IFITM-3-HA were obtained by Stephen Elledge (Brass et al., 2009). pQCXIP-IFITM3-HA rs12252, Δ1-21 and Y^20^A were generated by cloning corresponding gblocks (Integrated DNA Technologies) into pQXCIP using the *NotI* and *AgeI* restriction sites. Retroviral particles were generated by lipofection of HEK293T cells with pQCXIP-IFITM-HA constructs, and plasmids encoding MLV gag pol (Bartosch, Dubuisson et al., 2003) and pCMV-VSV-G (Stewart, Dykxhoorn et al., 2003). For production of lentiviral pseudotypes expressing luciferase, HEK293T cells were lipofected with pCSII-EF-luciferase (Agarwal, Nikolai et al., 2006), pCMV DR8.91 (Zufferey, Nagy et al., 1997) and a plasmid encoding indicated viral glycoprotein, MLV gp and Ebola gp (Chan, Speck et al., 2000). CHIKV glycoprotein mutations were introduced into the pIRES2-EGFP-CHIKV E3-E1 (Weber, Berberich et al., 2017), S27 isolate-based plasmid via site-directed mutagenesis using the QuikChange II Site-Directed Mutagenesis Kit (Agilent Technologies). The following mutations were introduced: E1(A^226^V); E1(A^226^V/ M^269^V/), E2(K^252^Q): sl1; E1(K^211^N/A^226^V), E2(V^222^I): sl2; E1(A^226^V/M^269^V), E3(S^18^F): sl3A; E1(A^226^V/M^269^V), E2(R^198^Q), E3(S^18^F): sl3B; E1(A^226^V/M^269^V), E2(L^210^Q): sl4. pCHIKrep1 EGFP, encoding CHIKV non-structural proteins 1-4 was kindly provided by Gorben Pijlman (Fros et al., 2010).

### Viruses

CHIKV was produced by electroporation of *in vitro* transcribed RNA derived from the molecular clone pCHIK-LR2006-OPY-5’EGFP (Tsetsarkin et al., 2007) into BHK-21 cells. Two days later, supernatant was filtered, and viral titers were determined by titration on HEK293T cells. *In vitro* transcribed full-length CHIKV mRNA and mRNA encoding single viral proteins was transfected into target cells with the *Trans*IT mRNA kit (Mirus). MAYV (strain TRVL15537) was passaged on Vero cells. HSV-1 stocks were prepared as previously described (Grosche *et al.*, 2020). Briefly, almost confluent BHK cells were infected at an MOI of 0.01 PFU/cell for 3 days with the BAC-derived strain HSV1(17^+^)Lox-_pMCMV_GFP, which expresses GFP under the control of the major immediate-early promoter of murine cytomegalovirus (Snijder, Sacher et al., 2012). The culture medium was collected, cells and debris were sedimented, and HSV-1 particles were harvested by high-speed centrifugation. The resulting virus pellets were resuspended in MNT buffer. Single-use stocks were aliquoted and kept at −80°C. IAV (strain A/PuertoRico/8/34 H1N1) was generated by an eight-plasmid rescue system kindly provided by Richard Webby (St. Jude Children’s Research Hospital, Memphis, TN), using transfection of HEK293T cells and subsequent infection of MDCK cells to generate viral progeny (Hoffmann, Krauss et al., 2002).

### Immunoblotting

Cells were lysed with M-PER Mammalian Protein Extraction Reagent (Pierce) and processed according to the recommended protocol. The lysate was mixed with Laemmli buffer and boiled for ten minutes at 95°C. Proteins were run on a 10 % SDS-PAGE and immobilized on a nitrocellulose membrane (GE Healthcare) using the Trans-Blot Turbo system (BioRad). Blocked membranes were incubated with the following antibodies: mouse anti-MAPK (clone D2, Santa Cruz Technologies, 1:1000), mouse anti-HA (clone 16B12, BioLegend, 1:1000), mouse anti-ISG15 (clone F-9, Santa Cruz Technologies, 1:500), mouse anti-IFITM1 (clone 5B5E2, Proteintech, 1:5000), mouse anti-IFITM2 (clone 3D5F7, Proteintech, 1:5000), rabbit anti-IFITM3 (Cat. nr. AP1153a, Abcepta, 1:500), rabbit anti-CHIKV antiserum (IBT Bioservices, 1:10,000) or mouse anti-p24 (ExBio, 1:1000). Secondary antibodies conjugated to Alexa680/800 fluorescent dyes were used for detection and quantification by Odyssey Infrared Imaging System (LI-COR Biosciences).

### Flow cytometry

Cells were fixed with 4 % PFA (Carl Roth) and permeabilized in 0.1 % Triton X-100 (Thermo Scientific) in PBS prior to immunostaining, if not stated otherwise. Cells were immunostained with the following antibodies: rabbit anti-IFITM3 (Cat. nr. AP1153a, Abcepta, 1:500) and mouse anti-HA (clone 16B12, BioLegend, 1:1400). Secondary antibodies conjugated to Alexa® Fluor 488 or 647 (Invitrogen, 1:1500) were used for detection. Flow cytometry analysis was performed using FACS Calibur or FACS Accuri with BD CellQuest Pro 4.0.2 Software (BD Pharmingen) and FlowJo V10 Software (FlowJo).

### Immunofluorescence Microscopy

Cells were grown in μ-slide 8 wells (Ibidi). Cells were fixed with 4 % PFA and permeabilized with 0.5 % Triton X-100 in PBS. Immunostaining was performed with primary antibodies for one hour at room temperature for HA (1:1000, BioLegend) or rabbit anti-IFITM3 (Cat. nr. AP1153a, Abcepta, 1:500) and secondary antibodies conjugated to Alexa Fluor® 488 and 647 (1:1000, Invitrogen) for one hour at room temperature. Nuclear DNA was stained with 2.5 μg/mL DAPI (Invitrogen) for five minutes at room temperature. Microscopic analysis was performed using a Nikon Ti-E microscope equipped with a Yokogawa CSU-X1 spinning-disc and an Andor DU-888 camera. ImageJ was used to prepare microscopy images and for quantification of signal intensity of the immunostaining.

#### Quantitative RT-PCR

Cells were lysed and RNA extracted using the Promega Maxwell 16 with the LEV simplyRNA tissue. cDNA was prepared using dNTPs (Thermo Fisher), random hexamers (Jena Bioscience) and M-MuLV reverse transcriptase (NEB) with buffer. Quantification of relative cellular mRNA levels was performed with the 7500 Fast Real-Time PCR System (Applied Biosystems) in technical triplicates using Taq-Man PCR technology with the following Taqman probes and primers (Thermo Fisher): human *IFITM3* (assay ID Hs03057129_s1), *IFIT1* (assay ID Hs01911452_s1), *RNASEP* (#4316849). Influenza virus RNA replication was assessed by quantifying HA mRNA levels with forward primer CAGATGCAGACACAATATGT and reverse primer TAGTGGGGCTATTCCTTTTA. Relative expression was calculated with the ◻◻CT method (Livak KJ et al Methods 2001), using *RNASEP* or *GAPDH* mRNA as reference.

#### Data Presentation and Statistical Analysis

If not otherwise stated, bars and symbols show the arithmetic mean of indicated amount of repetitions. Error bars indicate S.D. from at least three or S.E.M. from the indicated amount of individual experiments. Statistical analysis was performed with GraphPad Prism 8.3.0 using two-tailed unpaired Student’s t-tests. *P* values <0.05 were considered significant (*), <0.01 very significant (**), <0,001 extremely significant (***); n.s. = not significant (≥0.05).

## ACKNOWLEDGEMENTS

We thank Oliver Dittrich-Breiholz and the Transcriptomics Facility from Hanover Medical School, and Victor Tarabykin for granting access to the Step One Plus Real Time PCR System and the ABI7500 Real Time PCR System, respectively. We thank Hildegard Schilling for technical support with culture and typing of gene-edited cell lines. We thank Thomas Pietschmann and Christian Drosten for constant support.

This work was supported by funding from the Deutsche Forschungsgemeinschaft (DFG) for Germany’s Excellence Strategy, EXC 2155, project number 390874280 and SFB 900 – 158989968 (TPC2) awarded to B.S.; by grant GO2153/3-1 to C.G within DFG German/African Cooperation Projects in Infectiology, by the Impulse and Networking Fund of the Helmholtz Association through the HGF-EU partnering grant PIE-008 to C.G., funding of the Helmholtz Center for Infection Research (HZI) and of Berlin Institute of Health (BIH) to C.G.

## AUTHOR CONTRIBUTIONS

S.F., T.Z., F.P. and C.G. conceived and designed the experiments; S.F., T.Z., F.P., C.S., S.D., C.F., S.S., F.C., K.D. performed the experiments, S.F., T.Z., F.P., C.S., S.D., S.S., F.C. analyzed the data.; S.F. and C.G. drafted the manuscript.; S.F., F.P., K.D., B.S. and C.G. reviewed and edited the manuscript.; G.H., B.S., F.P., G.S., J.F.D., and C.G. supervised the research.

## SUPPLEMENTAL FIGURE LEGENDS

**Figure S1.**
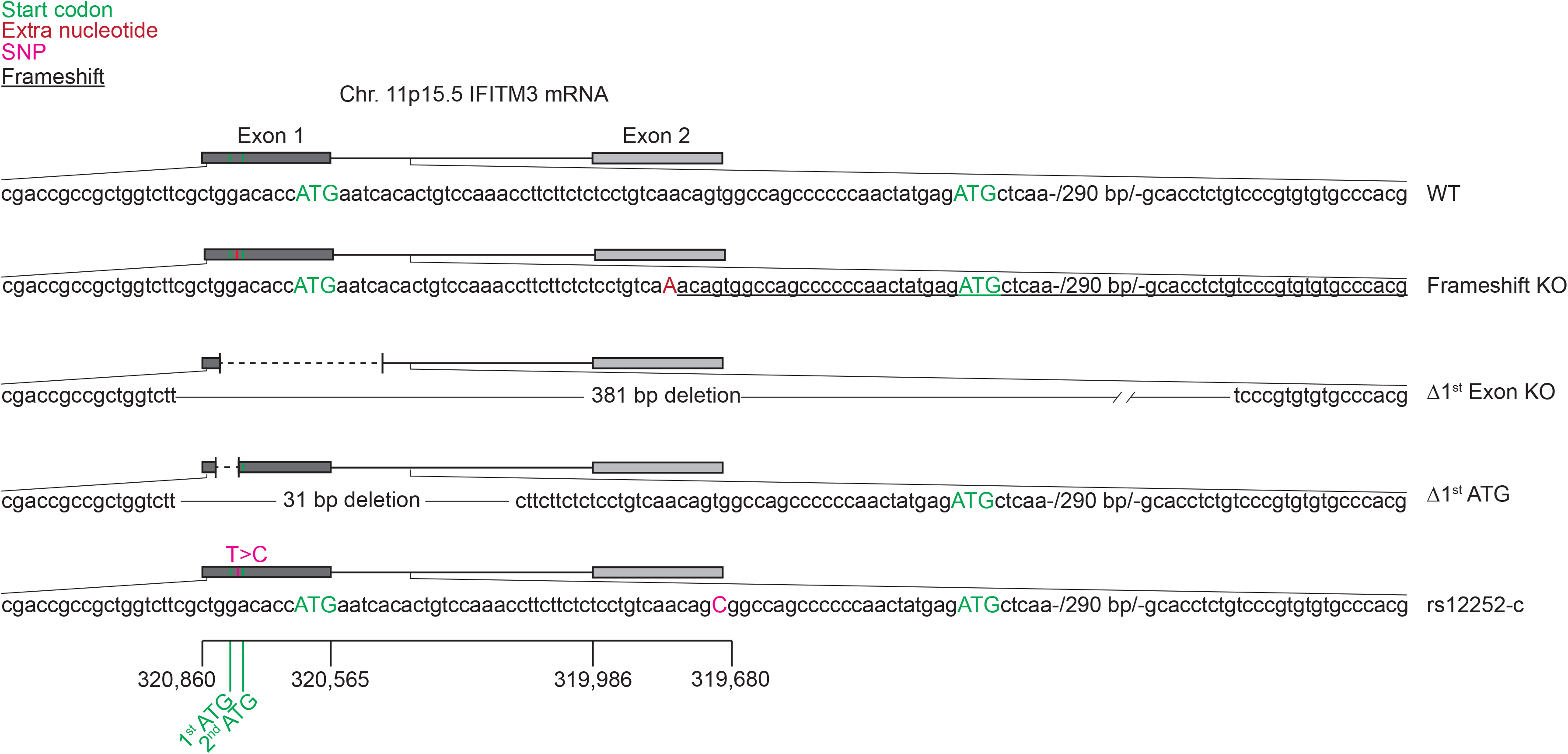
Schematic overview of *IFITM3* variants analyzed in this study

**Figure S2.**
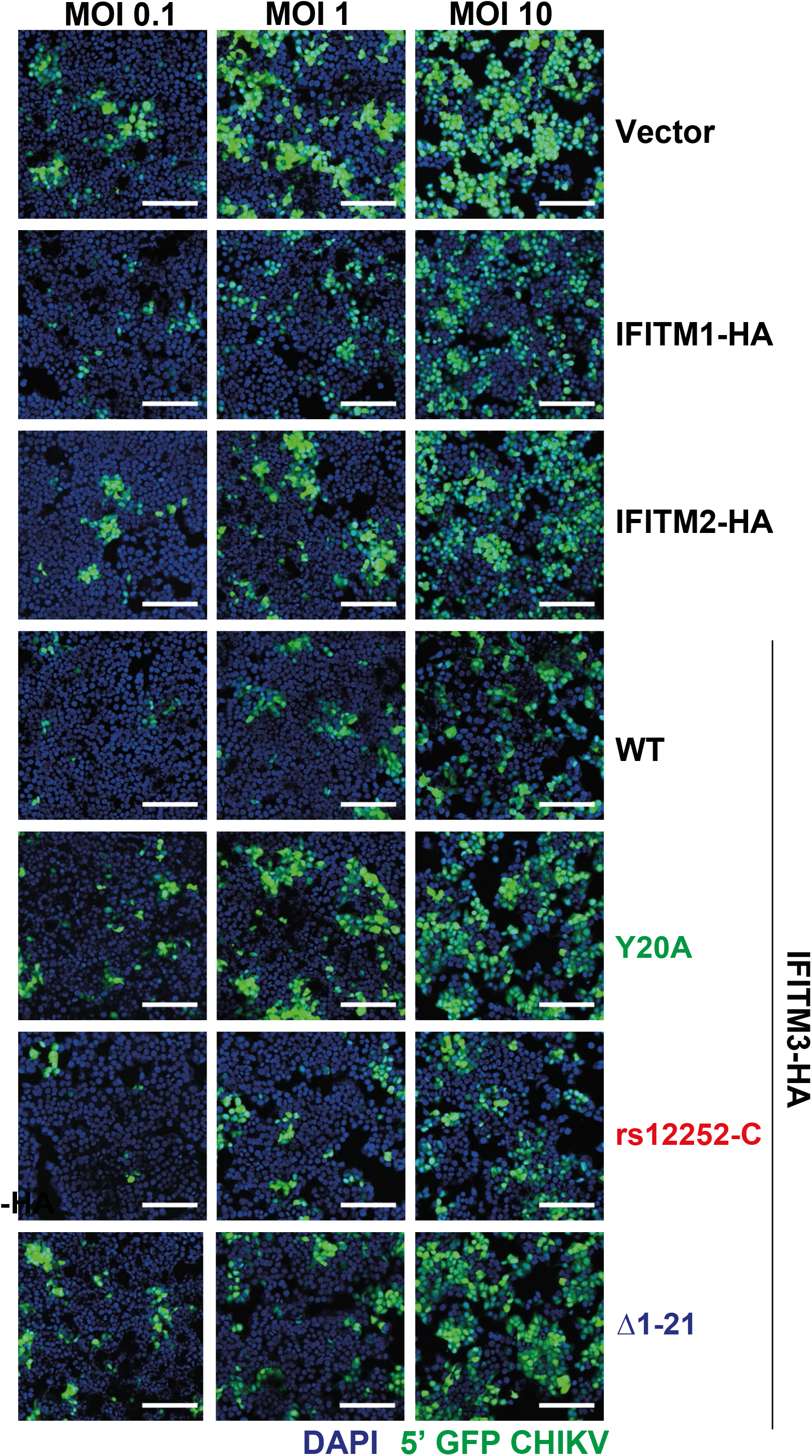
IFITM-HA expressing cells were infected with EGFP-CHIKV of increasing MOIs for 24 hours, and cells were analyzed by fluorescence microscopy.

## REFERENCES

Agarwal S, Nikolai B, Yamaguchi T, Lech P, Somia NV (2006) Construction and use of retroviral vectors encoding the toxic gene barnase. Mol Ther 14: 555–63

Akhrymuk I, Kulemzin SV, Frolova EI (2012) Evasion of the innate immune response: the Old World alphavirus nsP2 protein induces rapid degradation of Rpb1, a catalytic subunit of RNA polymerase II. J Virol 86: 7180–91

Appourchaux R, Delpeuch M, Zhong L, Burlaud-Gaillard J, Tartour K, Savidis G, Brass A, Etienne L, Roingeard P, Cimarelli A (2019) Functional Mapping of Regions Involved in the Negative Imprinting of Virion Particle Infectivity and in Target Cell Protection by Interferon-Induced Transmembrane Protein 3 against HIV-1. J Virol 93

Bailey CC, Zhong G, Huang IC, Farzan M (2014) IFITM-Family Proteins: The Cell's First Line of Antiviral Defense. Annu Rev Virol 1: 261–283

Bartosch B, Dubuisson J, Cosset FL (2003) Infectious hepatitis C virus pseudo-particles containing functional E1-E2 envelope protein complexes. J Exp Med 197: 633–42

Brass AL, Huang IC, Benita Y, John SP, Krishnan MN, Feeley EM, Ryan BJ, Weyer JL, van der Weyden L, Fikrig E, Adams DJ, Xavier RJ, Farzan M, Elledge SJ (2009) The IFITM proteins mediate cellular resistance to influenza A H1N1 virus, West Nile virus, and dengue virus. Cell 139: 1243–54

Carter TC, Hebbring SJ, Liu J, Mosley JD, Shaffer CM, Ivacic LC, Kopitzke S, Stefanski EL, Strenn R, Sundaram ME, Meece J, Brilliant MH, Ferdinands JM, Belongia EA (2018) Pilot screening study of targeted genetic polymorphisms for association with seasonal influenza hospital admission. J Med Virol 90: 436–446

Chan SY, Speck RF, Ma MC, Goldsmith MA (2000) Distinct mechanisms of entry by envelope glycoproteins of Marburg and Ebola (Zaire) viruses. J Virol 74: 4933–7

Chesarino NM, McMichael TM, Hach JC, Yount JS (2014) Phosphorylation of the antiviral protein interferon-inducible transmembrane protein 3 (IFITM3) dually regulates its endocytosis and ubiquitination. J Biol Chem 289: 11986–92

Chesarino NM, McMichael TM, Yount JS (2015) E3 Ubiquitin Ligase NEDD4 Promotes Influenza Virus Infection by Decreasing Levels of the Antiviral Protein IFITM3. PLoS Pathog 11: e1005095

Compton AA, Bruel T, Porrot F, Mallet A, Sachse M, Euvrard M, Liang C, Casartelli N, Schwartz O (2014) IFITM proteins incorporated into HIV-1 virions impair viral fusion and spread. Cell Host Microbe 16: 736–47

Compton AA, Roy N, Porrot F, Billet A, Casartelli N, Yount JS, Liang C, Schwartz O (2016) Natural mutations in IFITM3 modulate post-translational regulation and toggle antiviral specificity. EMBO Rep 17: 1657–1671

David S, Correia V, Antunes L, Faria R, Ferrao J, Faustino P, Nunes B, Maltez F, Lavinha J, Rebelo de Andrade H (2018) Population genetics of IFITM3 in Portugal and Central Africa reveals a potential modifier of influenza severity. Immunogenetics 70: 169–177

Desai TM, Marin M, Chin CR, Savidis G, Brass AL, Melikyan GB (2014) IFITM3 restricts influenza A virus entry by blocking the formation of fusion pores following virus-endosome hemifusion. PLoS Pathog 10: e1004048

Everitt AR, Clare S, Pertel T, John SP, Wash RS, Smith SE, Chin CR, Feeley EM, Sims JS, Adams DJ, Wise HM, Kane L, Goulding D, Digard P, Anttila V, Baillie JK, Walsh TS, Hume DA, Palotie A, Xue Y et al. (2012) IFITM3 restricts the morbidity and mortality associated with influenza. Nature 484: 519–23

Feeley EM, Sims JS, John SP, Chin CR, Pertel T, Chen L-M, Gaiha GD, Ryan BJ, Donis RO, Elledge SJ, Brass AL (2011) IFITM3 Inhibits Influenza A Virus Infection by Preventing Cytosolic Entry. PLOS Pathogens 7: e1002337

Fros JJ, Liu WJ, Prow NA, Geertsema C, Ligtenberg M, Vanlandingham DL, Schnettler E, Vlak JM, Suhrbier A, Khromykh AA, Pijlman GP (2010) Chikungunya virus nonstructural protein 2 inhibits type I/II interferon-stimulated JAK-STAT signaling. J Virol 84: 10877–87

Gorman MJ, Poddar S, Farzan M, Diamond MS (2016) The Interferon-Stimulated Gene Ifitm3 Restricts West Nile Virus Infection and Pathogenesis. J Virol 90: 8212–25

Hahon N, Zimmerman WD (1970) Chikungunya virus infection of cell monolayers by cell-to-cell and extracellular transmission. Appl Microbiol 19: 389–91

Hoffmann E, Krauss S, Perez D, Webby R, Webster RG (2002) Eight-plasmid system for rapid generation of influenza virus vaccines. Vaccine 20: 3165–70

Jia R, Pan Q, Ding S, Rong L, Liu SL, Geng Y, Qiao W, Liang C (2012) The N-terminal region of IFITM3 modulates its antiviral activity by regulating IFITM3 cellular localization. J Virol 86: 13697–707

Jia R, Xu F, Qian J, Yao Y, Miao C, Zheng YM, Liu SL, Guo F, Geng Y, Qiao W, Liang C (2014) Identification of an endocytic signal essential for the antiviral action of IFITM3. Cell Microbiol 16: 1080–93

John SP, Chin CR, Perreira JM, Feeley EM, Aker AM, Savidis G, Smith SE, Elia AE, Everitt AR, Vora M, Pertel T, Elledge SJ, Kellam P, Brass AL (2013) The CD225 domain of IFITM3 is required for both IFITM protein association and inhibition of influenza A virus and dengue virus replication. J Virol 87: 7837–52

Jolly C, Booth NJ, Neil SJ (2010) Cell-cell spread of human immunodeficiency virus type 1 overcomes tetherin/BST-2-mediated restriction in T cells. J Virol 84: 12185–99

Lee CY, Kam YW, Fric J, Malleret B, Koh EG, Prakash C, Huang W, Lee WW, Lin C, Lin RT, Renia L, Wang CI, Ng LF, Warter L (2011) Chikungunya virus neutralization antigens and direct cell-to-cell transmission are revealed by human antibody-escape mutants. PLoS Pathog 7: e1002390

Li C, Du S, Tian M, Wang Y, Bai J, Tan P, Liu W, Yin R, Wang M, Jiang Y, Li Y, Zhu N, Zhu Y, Li T, Wu S, Jin N, He F (2018) The Host Restriction Factor Interferon-Inducible Transmembrane Protein 3 Inhibits Vaccinia Virus Infection. Front Immunol 9: 228

Li C, Zheng H, Wang Y, Dong W, Liu Y, Zhang L, Zhang Y (2019) Antiviral Role of IFITM Proteins in Classical Swine Fever Virus Infection. Viruses 11

Li K, Markosyan RM, Zheng YM, Golfetto O, Bungart B, Li M, Ding S, He Y, Liang C, Lee JC, Gratton E, Cohen FS, Liu SL (2013) IFITM proteins restrict viral membrane hemifusion. PLoS Pathog 9: e1003124

Liao Y, Goraya MU, Yuan X, Zhang B, Chiu SH, Chen JL (2019) Functional Involvement of Interferon-Inducible Transmembrane Proteins in Antiviral Immunity. Front Microbiol 10: 1097

Liberatore RA, Mastrocola EJ, Powell C, Bieniasz PD (2017) Tetherin Inhibits Cell-Free Virus Dissemination and Retards Murine Leukemia Virus Pathogenesis. J Virol 91

Lopez-Rodriguez M, Herrera-Ramos E, Sole-Violan J, Ruiz-Hernandez JJ, Borderias L, Horcajada JP, Lerma-Chippirraz E, Rajas O, Briones M, Perez-Gonzalez MC, Garcia-Bello MA, Lopez-Granados E, Rodriguez de Castro F, Rodriguez-Gallego C (2016) IFITM3 and severe influenza virus infection. No evidence of genetic association. Eur J Clin Microbiol Infect Dis 35: 1811–1817

Lu J, Pan Q, Rong L, He W, Liu SL, Liang C (2011) The IFITM proteins inhibit HIV-1 infection. J Virol 85: 2126–37

Makvandi-Nejad S, Laurenson-Schafer H, Wang L, Wellington D, Zhao Y, Jin B, Qin L, Kite K, Moghadam HK, Song C, Clark K, Hublitz P, Townsend AR, Wu H, McMichael AJ, Zhang Y, Dong T (2018) Lack of Truncated IFITM3 Transcripts in Cells Homozygous for the rs12252-C Variant That is Associated With Severe Influenza Infection. J Infect Dis 217: 257–262

Meertens L, Hafirassou ML, Couderc T, Bonnet-Madin L, Kril V, Kummerer BM, Labeau A, Brugier A, Simon-Loriere E, Burlaud-Gaillard J, Doyen C, Pezzi L, Goupil T, Rafasse S, Vidalain PO, Bertrand-Legout A, Gueneau L, Juntas-Morales R, Ben Yaou R, Bonne G et al. (2019) FHL1 is a major host factor for chikungunya virus infection. Nature 574: 259–263

Mills TC, Rautanen A, Elliott KS, Parks T, Naranbhai V, Ieven MM, Butler CC, Little P, Verheij T, Garrard CS, Hinds C, Goossens H, Chapman S, Hill AV (2014) IFITM3 and susceptibility to respiratory viral infections in the community. J Infect Dis 209: 1028–31

Narayana SK, Helbig KJ, McCartney EM, Eyre NS, Bull RA, Eltahla A, Lloyd AR, Beard MR (2015) The Interferon-induced Transmembrane Proteins, IFITM1, IFITM2, and IFITM3 Inhibit Hepatitis C Virus Entry. J Biol Chem 290: 25946–59

Pan Y, Yang P, Dong T, Zhang Y, Shi W, Peng X, Cui S, Zhang D, Lu G, Liu Y, Wu S, Wang Q (2017) IFITM3 Rs12252-C Variant Increases Potential Risk for Severe Influenza Virus Infection in Chinese Population. Frontiers in cellular and infection microbiology 7: 294

Perreira JM, Chin CR, Feeley EM, Brass AL (2013) IFITMs restrict the replication of multiple pathogenic viruses. Journal of molecular biology 425: 4937–4955

Poddar S, Hyde JL, Gorman MJ, Farzan M, Diamond MS (2016) The Interferon-Stimulated Gene IFITM3 Restricts Infection and Pathogenesis of Arthritogenic and Encephalitic Alphaviruses. J Virol 90: 8780–94

Puigdomenech I, Casartelli N, Porrot F, Schwartz O (2013) SAMHD1 restricts HIV-1 cell-to-cell transmission and limits immune detection in monocyte-derived dendritic cells. J Virol 87: 2846–56

Randolph AG, Yip WK, Allen EK, Rosenberger CM, Agan AA, Ash SA, Zhang Y, Bhangale TR, Finkelstein D, Cvijanovich NZ, Mourani PM, Hall MW, Su HC, Thomas PG, Pediatric Acute Lung I, Sepsis Investigators Network Pediatric Influenza I, Pediatric Acute Lung I, Sepsis Investigators Network Pediatric Influenza I (2017) Evaluation of IFITM3 rs12252 Association With Severe Pediatric Influenza Infection. J Infect Dis 216: 14–21

Rausalu K, Utt A, Quirin T, Varghese FS, Zusinaite E, Das PK, Ahola T, Merits A (2016) Chikungunya virus infectivity, RNA replication and non-structural polyprotein processing depend on the nsP2 protease's active site cysteine residue. Sci Rep 6: 37124

Richardson MW, Carroll RG, Stremlau M, Korokhov N, Humeau LM, Silvestri G, Sodroski J, Riley JL (2008) Mode of transmission affects the sensitivity of human immunodeficiency virus type 1 to restriction by rhesus TRIM5alpha. J Virol 82: 11117–28

Savidis G, Perreira JM, Portmann JM, Meraner P, Guo Z, Green S, Brass AL (2016) The IFITMs Inhibit Zika Virus Replication. Cell Rep 15: 2323–30

Schoggins JW, Wilson SJ, Panis M, Murphy MY, Jones CT, Bieniasz P, Rice CM (2011) A diverse range of gene products are effectors of the type I interferon antiviral response. Nature 472: 481–5

Snijder B, Sacher R, Ramo P, Liberali P, Mench K, Wolfrum N, Burleigh L, Scott CC, Verheije MH, Mercer J, Moese S, Heger T, Theusner K, Jurgeit A, Lamparter D, Balistreri G, Schelhaas M, De Haan CA, Marjomaki V, Hyypia T et al. (2012) Single-cell analysis of population context advances RNAi screening at multiple levels. Mol Syst Biol 8: 579

Spence JS, He R, Hoffmann HH, Das T, Thinon E, Rice CM, Peng T, Chandran K, Hang HC (2019) IFITM3 directly engages and shuttles incoming virus particles to lysosomes. Nat Chem Biol 15: 259–268

Stewart SA, Dykxhoorn DM, Palliser D, Mizuno H, Yu EY, An DS, Sabatini DM, Chen IS, Hahn WC, Sharp PA, Weinberg RA, Novina CD (2003) Lentivirus-delivered stable gene silencing by RNAi in primary cells. RNA 9: 493–501

Suddala KC, Lee CC, Meraner P, Marin M, Markosyan RM, Desai TM, Cohen FS, Brass AL, Melikyan GB (2019) Interferon-induced transmembrane protein 3 blocks fusion of sensitive but not resistant viruses by partitioning into virus-carrying endosomes. PLoS Pathog 15: e1007532

Tartour K, Appourchaux R, Gaillard J, Nguyen XN, Durand S, Turpin J, Beaumont E, Roch E, Berger G, Mahieux R, Brand D, Roingeard P, Cimarelli A (2014) IFITM proteins are incorporated onto HIV-1 virion particles and negatively imprint their infectivity. Retrovirology 11: 103

Timpe JM, Stamataki Z, Jennings A, Hu K, Farquhar MJ, Harris HJ, Schwarz A, Desombere I, Roels GL, Balfe P, McKeating JA (2008) Hepatitis C virus cell-cell transmission in hepatoma cells in the presence of neutralizing antibodies. Hepatology 47: 17–24

Tsetsarkin KA, Chen R, Yun R, Rossi SL, Plante KS, Guerbois M, Forrester N, Perng GC, Sreekumar E, Leal G, Huang J, Mukhopadhyay S, Weaver SC (2014) Multi-peaked adaptive landscape for chikungunya virus evolution predicts continued fitness optimization in Aedes albopictus mosquitoes. Nat Commun 5: 4084

Tsetsarkin KA, Vanlandingham DL, McGee CE, Higgs S (2007) A Single Mutation in Chikungunya Virus Affects Vector Specificity and Epidemic Potential. PLoS Pathogens 3: e201

Vendrame D, Sourisseau M, Perrin V, Schwartz O, Mammano F (2009) Partial inhibition of human immunodeficiency virus replication by type I interferons: impact of cell-to-cell viral transfer. J Virol 83: 10527–37

Weber C, Berberich E, von Rhein C, Henss L, Hildt E, Schnierle BS (2017) Identification of Functional Determinants in the Chikungunya Virus E2 Protein. PLoS Negl Trop Dis 11: e0005318

Weidner JM, Jiang D, Pan XB, Chang J, Block TM, Guo JT (2010) Interferon-induced cell membrane proteins, IFITM3 and tetherin, inhibit vesicular stomatitis virus infection via distinct mechanisms. J Virol 84: 12646–57

Weston S, Czieso S, White IJ, Smith SE, Kellam P, Marsh M (2014) A membrane topology model for human interferon inducible transmembrane protein 1. PLoS One 9: e104341

Weston S, Czieso S, White IJ, Smith SE, Wash RS, Diaz-Soria C, Kellam P, Marsh M (2016) Alphavirus Restriction by IFITM Proteins. Traffic 17: 997–1013

White LK, Sali T, Alvarado D, Gatti E, Pierre P, Streblow D, Defilippis VR (2011) Chikungunya virus induces IPS-1-dependent innate immune activation and protein kinase R-independent translational shutoff. J Virol 85: 606–20

Williams DE, Wu WL, Grotefend CR, Radic V, Chung C, Chung YH, Farzan M, Huang IC (2014) IFITM3 polymorphism rs12252-C restricts influenza A viruses. PLoS One 9: e110096

Wrensch F, Karsten CB, Gnirß K, Hoffmann M, Lu K, Takada A, Winkler M, Simmons G, Pöhlmann S (2015) Interferon-Induced Transmembrane Protein-Mediated Inhibition of Host Cell Entry of Ebolaviruses. The Journal of infectious diseases 212 Suppl 2: S210–S218

Yount JS, Karssemeijer RA, Hang HC (2012) S-palmitoylation and ubiquitination differentially regulate interferon-induced transmembrane protein 3 (IFITM3)-mediated resistance to influenza virus. J Biol Chem 287: 19631–41

Zhang R, Kim AS, Fox JM, Nair S, Basore K, Klimstra WB, Rimkunas R, Fong RH, Lin H, Poddar S, Crowe JE, Jr., Doranz BJ, Fremont DH, Diamond MS (2018) Mxra8 is a receptor for multiple arthritogenic alphaviruses. Nature 557: 570–574

Zhang YH, Zhao Y, Li N, Peng YC, Giannoulatou E, Jin RH, Yan HP, Wu H, Liu JH, Liu N, Wang DY, Shu YL, Ho LP, Kellam P, McMichael A, Dong T (2013) Interferon-induced transmembrane protein-3 genetic variant rs12252-C is associated with severe influenza in Chinese individuals. Nat Commun 4: 1418

Zufferey R, Nagy D, Mandel RJ, Naldini L, Trono D (1997) Multiply attenuated lentiviral vector achieves efficient gene delivery in vivo. Nat Biotechnol 15: 871–5

